# Evidence of interplays between the vascular and nociceptive systems revealed by changes in capsaicin pain caused by limb position change

**DOI:** 10.1101/2024.01.16.575811

**Authors:** Arthur S. Courtin, Clara Knaepen, André Mouraux, Sabien Geraldine Antonia van Neerven

## Abstract

This experiment aimed at confirming our incidental observation that, when capsaicin is applied on the volar forearm, raising the arm to a vertical position leads to a dramatic increase in capsaicin-evoked pain and to explore possible underlying mechanisms.

Twenty healthy volunteers received a 2% capsaicin patch on one forearm and a vehicle patch on the other. Patches were kept in place for 60 minutes. The sensation caused by the patch was assessed repeatedly -in resting position and when the arm was raised vertically-before, during and after patch application. In addition, capsaicin-induced secondary hyperalgesia was assessed using mechanical pinprick stimuli. Half of the participants were seated upright while the other half were lying supine, to assess whether the effect of limb position was due to gravity.

After a few minutes of patch application, raising the capsaicin treated arm (but not the vehicle treated arm) led to a strong increase of the pain experienced at the patch. This effect of raising the arm did not differ between participants in the supine and seated groups and is therefore likely related to the position of the arm relative to the ground (gravity) rather than to the body. Mechanical secondary hyperalgesia and the arm raising effect were strongly decorrelated at the last time point after patch removal, indicating different underlying mechanisms.

Our results indicate that capsaicin-evoked pain can be strongly modulated by limb posture and that this effect may be caused by an interplay between vascular and nociceptive systems.

## 1. Introduction

When applied onto the human skin, capsaicin, a chemical naturally found in chili peppers, elicits a burning and painful sensation within a few minutes [14,28]. Capsaicin binds to the Transient Receptor Potential channel Vanilloid-1 (TRPV1), a ubiquitous heat sensitive nonselective cation channel, and shifts its activation threshold towards lower temperatures [12,37].

In the skin, TRPV1 is expressed by small diameter myelinated Aδ and unmyelinated C fibers, and by several non-neural cell types such as keratinocytes and schwann cells [1,12,15,18,27–29]. The burning sensation induced by topical capsaicin is thought to result from the reduced thermal activation threshold of TRPV1, leading to sustained activity in heat-sensitive nociceptors at baseline skin temperatures [28,37]. As some of these nociceptors are peptidergic fibers, their sustained activation also leads to the development of a flare in and around the area of capsaicin application [11,16].

Topical capsaicin also induces sensory changes that extend beyond the treated skin and involve more than heat signaling. For example, it induces dynamic mechanical allodynia and, with some delay relative to the capsaicin-induced burning sensation, a prolonged increase in sensitivity to mechanical pinprick stimuli delivered to the surrounding untreated skin [28,32]. This mechanical secondary hyperalgesia is thought to result from a heterosynaptic long-term potentiation (LTP) of dorsal horn neurons relaying inputs conveyed by mechanosensitive nociceptors, one of the first described instances of central sensitization [9,18].

In a previous study from our group during which the temporal evolution of sensory alterations caused by high-concentration topical capsaicin application in healthy volunteers was studied, we accidently observed that raising the capsaicin-treated arm from a neutral position to a vertical position dramatically increased burning pain intensity within a few seconds (observation not published) [27]. Raising the untreated contralateral arm did not elicit this increase in intensity.

This observation could be related to the report by Byas-Smith *et al.* that, after intradermal injection of capsaicin at the volar forearm, pain ratings increased markedly during arterial occlusion (using tourniquet constriction of the treated arm) and went back to their original values within a few seconds once blood flow was restored [10]. Indeed, raising the arm can be expected to lead to a reduction of blood flow in the arm, not unlike (mild) tourniquet occlusion. Interestingly, Byas-Smith *et al.* also reported (even though without showing data) that this “tourniquet allodynia” appeared only after pinprick hyperalgesia had developed, which could indicate a shared mechanism.

To further explore this matter, this preregistered study had several goals. First, to confirm our initial observation that simply holding the arm vertically markedly increases capsaicin-induced burning pain at the treated arm (i.e. the arm-raising effect). Second, to confirm that this arm-raising effect is related to the position of the arm relative to the ground (effect of gravitation on the vascular system) and not relative to the body. Third, to investigate whether the development of the arm-raising effect follows the time course of capsaicin-induced secondary hyperalgesia (manifested as an increase in pinprick hypersensitivity).

## 2. Methods

All experiments were conducted according to the latest version (October 2013) of the Declaration of Helsinki. Approval for the conduction of the experiment was obtained from the local ethical committee (Comité d’Ethique Hospitalo-Facultaire CUSL-UCLouvain, protocol 2019/15MAI/215 HPoC) before the recruitment of participants started. This study was pre-registered and the preregistration record as well as data and analysis code can be found on the corresponding OSF repository (DOI 10.17605/OSF.IO/XJ8BY).

### Participants

Twenty healthy young volunteers (age range: 18 – 30, median age: 22, 10 males, 10 females, 5 left-handed, 15 right-handed) were included. They all satisfied the following inclusion criteria: being between 18 and 30 years old; not using drugs on a regular basis (i.e. less than once a week; anticonception not included); not having a history of neurological or psychiatric condition; not having dermatological conditions or skin lesions on the forearms; not suffering of chronic pain or being in pain at the time of the experiment; not having a hobby that leads to over stimulation of the volar forearm (e.g. volleyball). All participants gave written informed consent before the beginning of the experiment.

### Capsaicin and vehicle patch application

Two patches were applied on each participant, one with capsaicin solution and the other with vehicle solution only. Capsaicin solution was obtained by diluting capsaicin powder (M2028, Sigma-Aldrich) in a vehicle solution consisting of 50% water and 50% ethanol absolute to a final concentration of 2% capsaicin. Each type of patch was obtained by dripping 1 mL of the corresponding solution on a 5x5 cm gauze pad (Sterilux ES, Hartmann) set on a 10 x 8.5 cm self-adhesive film (Opsite Flexifix, Smith and nephew).

**Figure 1:**
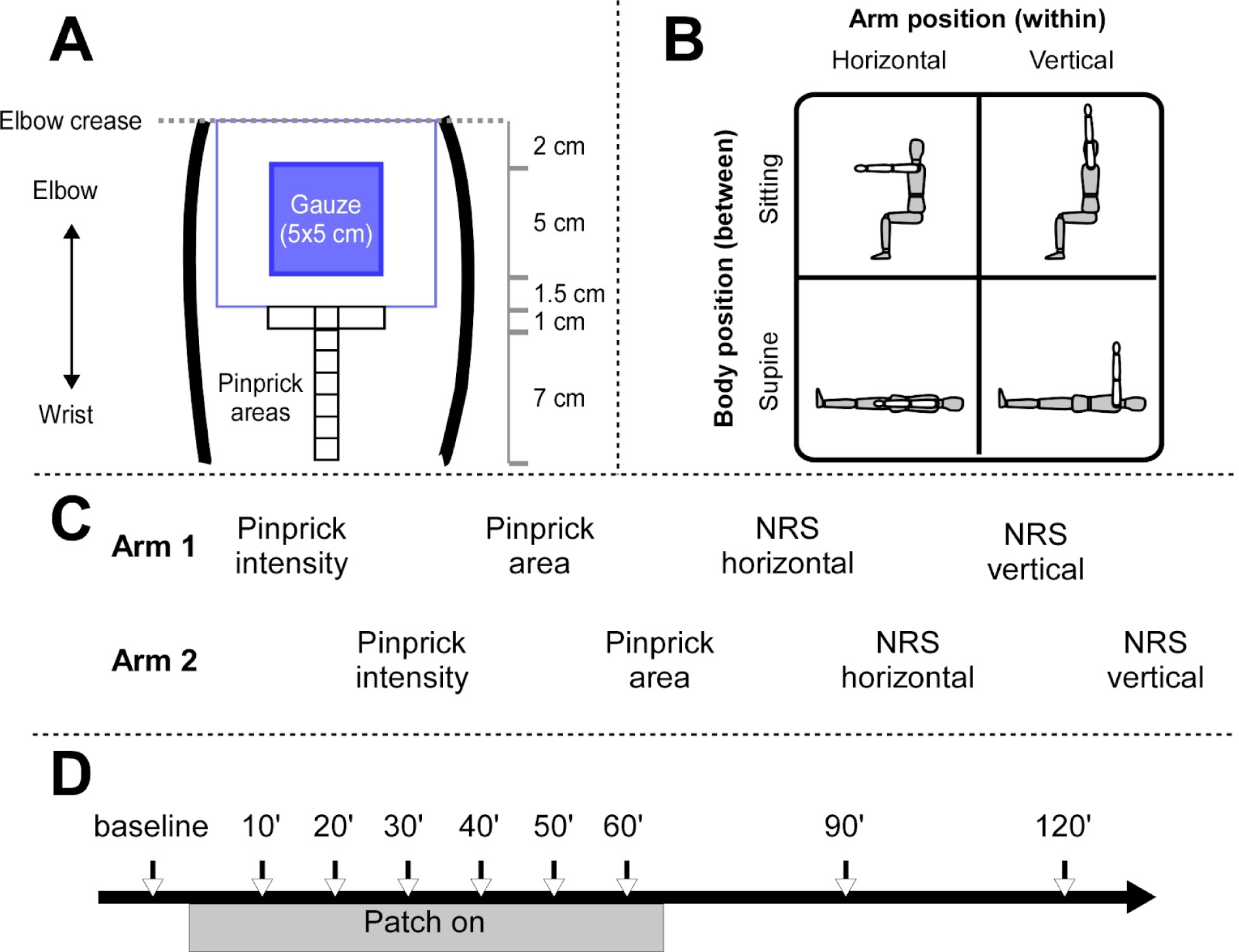
Experimental design. **A.** Placement of the patch and location of the pinprick stimulation areas. **B.** Representation of the levels of between subject factor “Body position” and within subject factor “Arm position”. **C**. Representation of one measurement set. **D**. Representation of the timeline of one experimental session, showing the timing of the different measurement sets relative to patch application.

### Assessment of sensations elicited by the patch

During the experiment, participants were repeatedly asked to report the intensity of the sensation elicited by the patches. To do so, they were asked to focus on the skin site corresponding to one of the patch for 20 s and to give a rating on a 0 to 100 numerical rating scale (NRS), where 0 corresponded to “*no sensation*” and 100 to “*the most intense sensation you can imagine*”. They were also asked to provide descriptors of the sensation elicited by the patch, in the form of epithet(s) from the following list and, if necessary, additional freely-chosen epithets: warm (*tiède*), hot (*chaud*), burning (*brûlant*), pricking (*piquant*), painful (*douloureux*), itchy (*démangeaison*), touch (*toucher*).

### Assessment of sensations elicited by the pinprick stimuli

To assess secondary mechanical hyperalgesia development, a 256 mN pinprick stimulator (The Pinprick, MRC Systems) was used. At each time point, three pinprick stimuli were delivered within a 1 x 5 cm area located adjacent to the distal edge of the self-adhesive film (Figure 1A) and the participant was asked to rate and provide qualifiers for each of these stimuli using the procedures described in the previous section. The target of the pinprick stimulator was pseudo-randomly changed after each stimulus to avoid sensitization due to repetitive stimulation of the same skin spot.

### Assessment of the spatial extent of secondary hyperalgesia

We also attempted to quantify the spatial extent of the secondary hyperalgesia. To do so, pinprick stimuli were delivered every cm on a 7 cm line starting at the distal edge of the self-adhesive film and following the axis of the forearm (Figure 1A). After each stimulus, participants were asked to indicate whether they felt the stimulus as clearly weaker (if stimuli were delivered from proximal to distal) or stronger (if stimuli were delivered from distal to proximal) than the previous one. At each time point, four such trains of stimuli were delivered on each forearm, two going from distal to proximal and two from proximal to distal. The length of the line connecting the self-adhesive film and the stimulation site at which the sensation was first reported as significantly stronger (weaker) for distal to proximal (proximal to distal) trains was recorded, as it should have corresponded to the edge of the pinprick allodynia area.

If valid, this measure should have resulted in a vast majority of 0 lengths for the vehicle patch and in matching lengths for opposite direction trains delivered next to the capsaicin patch. Whereas we observed this pattern in a few participants (usually people who previously participated in experiments involving secondary hyperalgesia), it was not the case for the majority. We therefore decided to not analyze these length data as what exactly was measured is unclear and likely variable across individuals.

### Experimental procedure

The experiment took place in a dimly lit and warm room. Before the beginning of the actual data collection, participants were explained and familiarized with the different measures. Then, participants had to practice the rating and description task on three 128 mN pinprick stimuli, three 256 mN pinprick stimuli, three 44°C stimuli and three 48°C stimuli, applied to their hand dorsum. Noxious heat stimuli lasted 5 s and were delivered with the T06 thermode of the TCSII (Peltier effect thermode, QST.Lab), roughly matching in size the area of the patch. Finally, the area on which the patches would be applied was marked on the forearms, using an ethanol resistant marker.

Actual data collection started with a first set of measurements. First, sensations elicited by pinprick stimuli were assessed as described previously for one forearm and then for the other. Second, the procedure described to quantify the spatial extent of secondary hyperalgesia was conducted for one forearm and then the other. Third, participants were asked to provide a rating for sensations at the site of the (future) patch, while keeping their arm in the horizontal position, for one forearm and then the other. Finally, this last step was repeated but, this time, participants had to raise their arm in a vertical position for 20 s and give a rating corresponding to the sensation experienced during the arm lift. The side with which the measurement set started (dominant *vs* non-dominant) was counterbalanced across participants.

Immediately after completion of the first measurement set, a capsaicin patch was applied on one of the participant’s volar forearms and a vehicle patch on the other. The patches were placed in such a way that the proximal edge of the gauze pad was located ∼2 cm away from the elbow crease (Figure 1A). The side to which the vehicle and capsaicin patches were applied (dominant *vs* non-dominant) was counterbalanced across participants.

To assess whether the position of the limb relative to the body (0° to ∼180° shoulder flexion) or relative to the ground (∼horizontal or ∼vertical) was driving the arm raising-effect, half of the participants were lying on their back (supine position) and half of the participants were sitting in a comfortable chair during the experiment with a high table in front of them to support and keep their arms horizontal (Figure 1B).

Six sets of measurements were taken while the patch was on, one every 10’ starting from 10’ after patch application (Figure 1C and D). After the end of the sixth measurement set (∼65’ post patch application), the patches were removed and the skin was gently cleaned with tepid water and soap, in order to remove capsaicin and vehicle solution residues. Two additional measurement sets were taken at 90’ and 120’ post patch application (roughly 25’ and 55’ post patch removal).

### Data analysis

Data analysis was conducted in R using the RStudio interface [3]. In addition to base R, functions from the *tidyverse* [38]*, ggeffects* [25] and *LmerTest* [24] packages were used.

To confirm that capsaicin led to ‘spontaneous’ burning sensations and that these sensations were increased during arm elevation, an ANOVA based on a linear mixed model -with a random intercept for each participant and fixed effect factors *time point* (9 levels), *arm position* (2 levels, horizontal vs vertical), *body position* (2 levels, supine vs sitting), *patch* (2 levels, vehicle vs capsaicin), and all their interactions-was used to analyze the patch ratings. Based on our hypotheses, we expected ratings for the vehicle patch to remain stable (and close to 0) over time and arm position, whereas the ratings from the capsaicin patch should increase over time (and maybe plateau or start decreasing at some point) and should be increased by arm elevation. This would correspond to a main effect of patch, and interactions between patch, time point, and arm position.

An additional, not pre-registered analysis of the probability of describing the patch stimuli with the descriptors “pricking” and/or “burning” and/or “painful” as a function of factors *time point* (9 levels), *arm position* (2 levels, horizontal vs vertical), and *patch* (2 levels, vehicle vs capsaicin) was performed, to check whether quantitative changes in the sensation (ratings) were matched with qualitative changes. For this analysis, an ANOVA based on a logistic mixed effects model with a random intercept for each participant was used. The amount of data (one value *per* factor combination) did not allow computation of interaction effects.

To confirm that capsaicin treatment led to the development of secondary mechanical hyperalgesia, an ANOVA based on a linear mixed model -with a random intercept for each participant and fixed effect factors *time point* (9 levels) and *patch* (2 levels, vehicle vs capsaicin), and all their interactions-was used to analyze the pinprick ratings. Based on our hypotheses, we expected to observe stable and low pinprick ratings on the forearm that received the vehicle patch and to observe increasing (over time) pinprick ratings on the arm that received the capsaicin patch (possibly followed by a plateau and decay). This would correspond to a main effect of patch and time point, as well as an interaction between these factors. In order to satisfy the distributional assumption of the model (normality of the residuals), the ratings had to be log-transformed.

An additional, not pre-registered analysis of the probability of describing the pinprick stimuli with the descriptors “pricking” and/or “burning” and/or “painful” as a function of factors *time point* (9 levels) and *patch* (2 levels, vehicle vs capsaicin) was performed, to check whether quantitative changes in the sensation (ratings) were matched with qualitative changes. For this analysis, an ANOVA based on a logistic mixed effects model with a random intercept for each participant was used.

As mentioned before, we decided not to analyze the spatial extent of pinprick hyperalgesia.

The pre-registered measure of temporal association between the arm-raising effect and mechanical secondary hyperalgesia proved impractical to compute due to the imbalance between pinprick ratings (3 *per* condition combinations) and the patch ratings (1 *per* conditions combinations). Instead of losing information by averaging pinprick ratings, we decided to use visual inspection of the marginal means (obtained from the previously described models) of the capsaicin patch rating in horizontal position (resting position), of the difference between the capsaicin patch ratings obtained in vertical and horizontal position (the “arm raising” effect), and of the difference between the pinprick ratings obtained from the vehicle and capsaicin arms (secondary hyperalgesia). For ease of comparison, these marginal means were normalized by dividing them (for each variable separately) by the largest marginal mean, so that each time course saturated at 1.

The alpha level was set to 0.05 and all plots show the marginal means and 95% confidence intervals fitted by the model.

## 3. Results

### Capsaicin treatment

Right after removal of the capsaicin patch, a clear flare response was visible at the treated volar forearm, extending several centimeters outside the area of the patch. At the vehicle treated arm, no flare was detectable upon visual inspection. Participants qualified the sensations elicited by the capsaicin patch as ‘warm’, ‘burning’, ‘pricking’, ‘itching’, or ‘painful’ (Figure 3). At the vehicle treated arm, the sensation of the patch on the skin was reported by a few participants and sometimes described as slightly ‘warm’ or ‘cold’ during the first time point after patch application.

**Figure 2:**
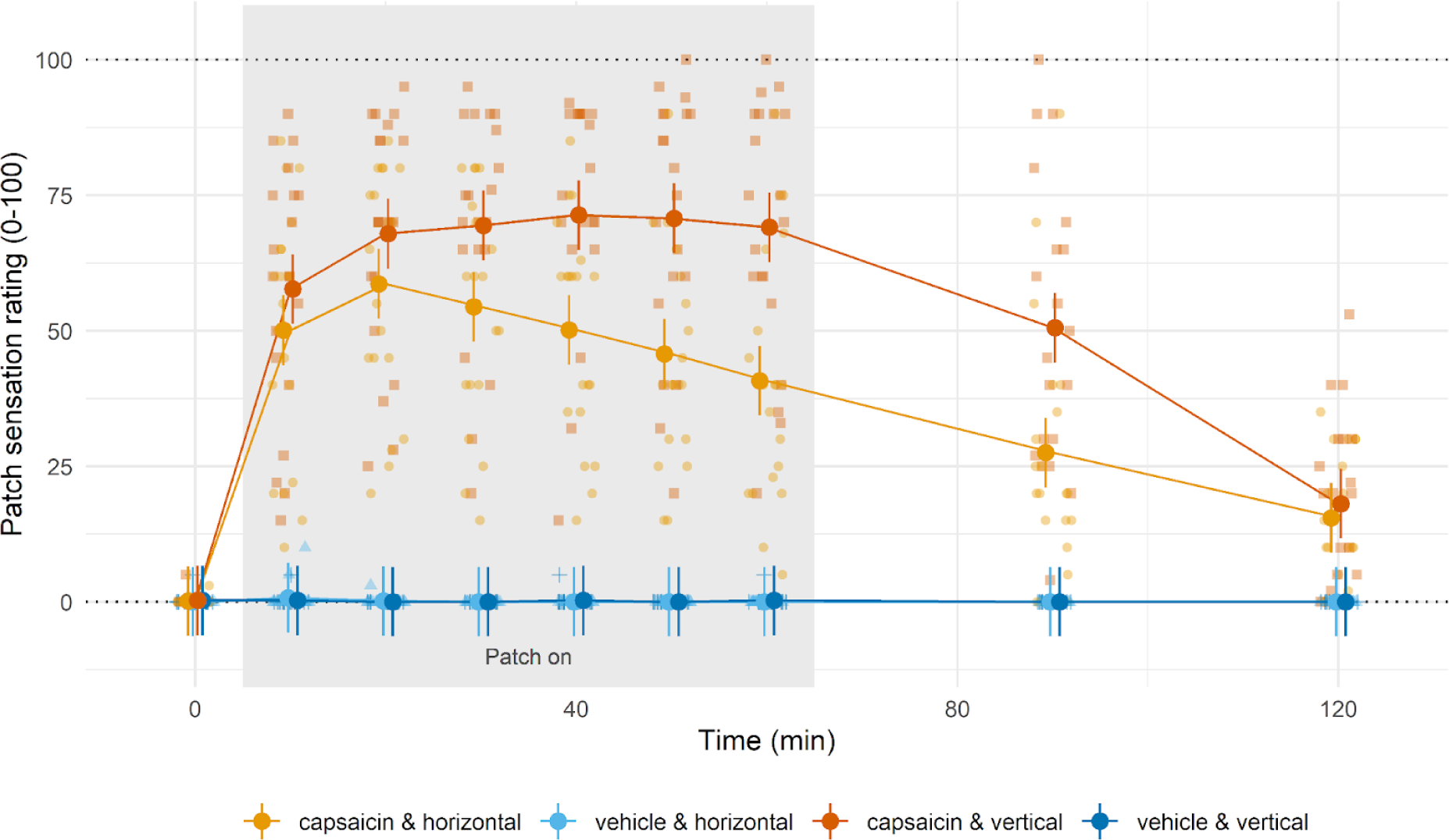
Perceived intensity of the sensations elicited by the patches, over time and across conditions. Individual ratings are represented as translucent dots in the background. The solid dots and vertical lines represent the marginal means and 95% confidence intervals fitted by the model while averaging over the levels of “body position” (as this factor had no significant effect on ratings). The solid lines connect the marginal means.

### Patch sensation

Sensations at the vehicle patch were mostly rated as 0, corresponding to no detectable sensation. This absence of patch evoked sensations remained stable over time and was not influenced by arm (Figure 2, light-blue and dark-blue lines) or body position. The capsaicin patch evoked strong ‘burning hot’ sensations (Figure 2 and 3, light-orange line) that quickly increased after patch application, plateaued around +20 minutes, and then slowly decayed (Table 1 and 2). Raising the arm while wearing the capsaicin patch produced a clear augmentation of these burning sensations (Figure 2 and 3, dark-orange line; Table 1 and 2).

Body position did not appear to influence the arm-raising effect (all factor combinations including “body position” had p>0.05 whereas all the other coefficients had p≤0.001), suggesting that it was the position of the arm relative to the ground (and not to the rest of the body) that drove the increase in sensation while raising the arm (Table 1).

**Figure 3:**
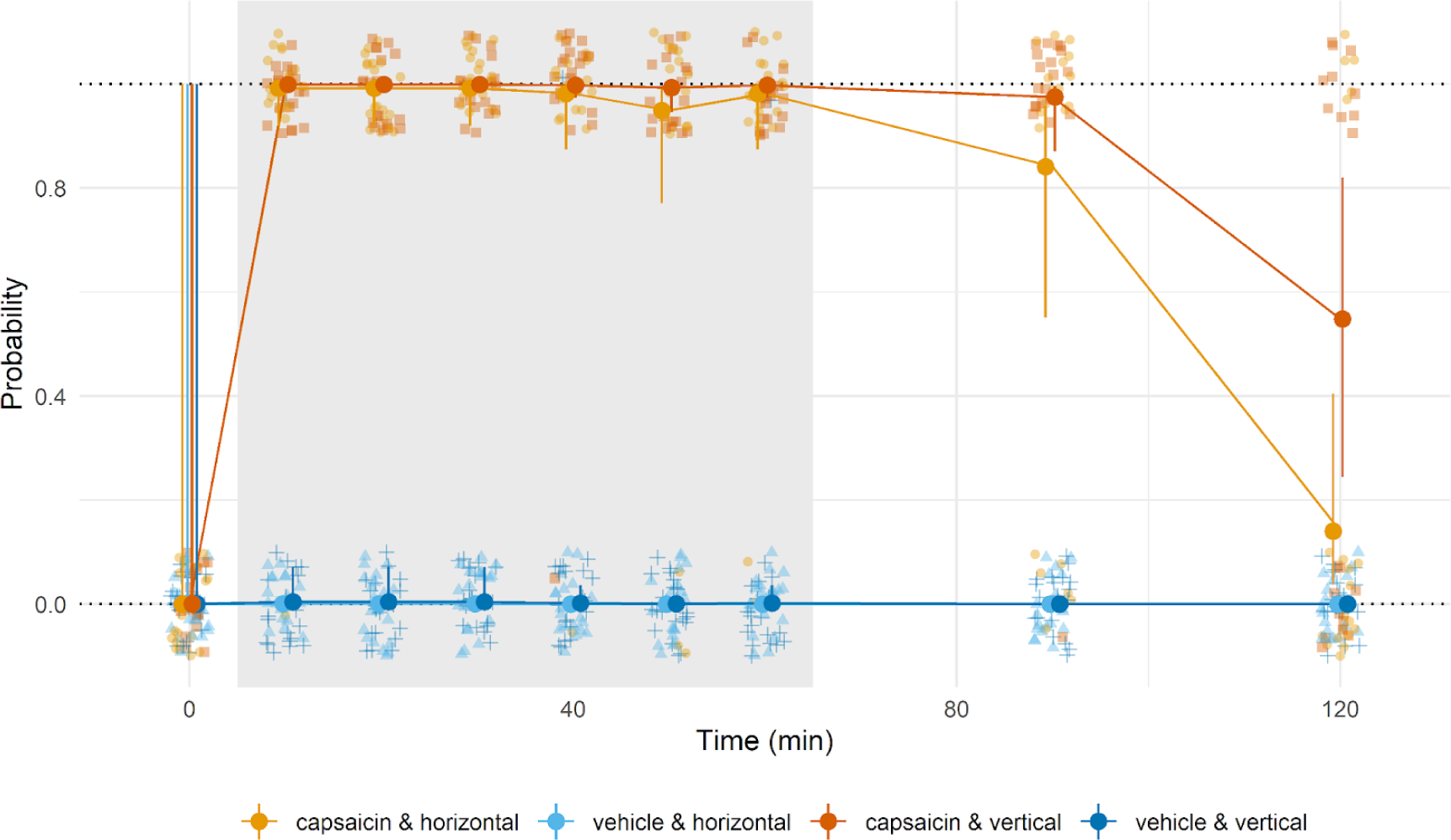
Qualitative assessment of sensations elicited by patch: probability of reporting a nociceptive descriptor (burning/pricking/painful). Individual responses are represented as translucent jittered dots in the background. The solid dots and vertical lines represent the marginal means and 95% confidence intervals fitted by the model. The solid lines connect the marginal means.

**Table 1:** Results of the patch ratings ANOVA.

| Factor | Sum Square | Mean Square | Num<br>DF | Den<br>DF | F value | Pr (>F) |
| --- | --- | --- | --- | --- | --- | --- |
| time point | 81275.894 | 10159.487 | 8 | 630 | 62.450 | <0.001 |
| arm position | 9709.381 | 9709.381 | 1 | 630 | 59.683 | <0.001 |
| body position | 20.522 | 20.522 | 1 | 18 | 0.126 | 0.727 |
| patch | 370101.144 | 370101.144 | 1 | 630 | 2275.000 | <0.001 |
| time point: arm position | 4273.694 | 534.212 | 8 | 630 | 3.284 | 0.001 |
| time point: body position | 215.950 | 26.994 | 8 | 630 | 0.166 | 0.995 |
| arm position: body position | 172.089 | 172.089 | 1 | 630 | 1.058 | 0.304 |
| time point: patch | 81035.444 | 10129.431 | 8 | 630 | 62.265 | <0.001 |
| arm position: patch | 9679.996 | 9679.996 | 1 | 630 | 59.503 | <0.001 |
| body position: patch | 264.023 | 264.023 | 1 | 630 | 1.623 | 0.203 |
| time point: arm position:<br>body position | 170.061 | 21.258 | 8 | 630 | 0.131 | 0.998 |
| time point: arm position:<br>patch | 4158.800 | 519.850 | 8 | 630 | 3.196 | 0.001 |
| time point: body position:<br>patch | 247.878 | 30.985 | 8 | 630 | 0.190 | 0.992 |
| arm position: body position:<br>patch | 168.200 | 168.200 | 1 | 630 | 1.034 | 0.310 |
| time point: arm position:<br>body position: patch | 179.300 | 22.412 | 8 | 630 | 0.138 | 0.997 |

**Table 2:** Results of the patch quality ANOVA.

| Factor | Chi Square | DF | Pr(>ChSq) |
| --- | --- | --- | --- |
| Patch | 47.297 | 1 | <0.001 |
| Time Point | 45.886 | 8 | <0.001 |
| Arm position | 11.997 | 1 | 0.001 |

**Figure 4:**
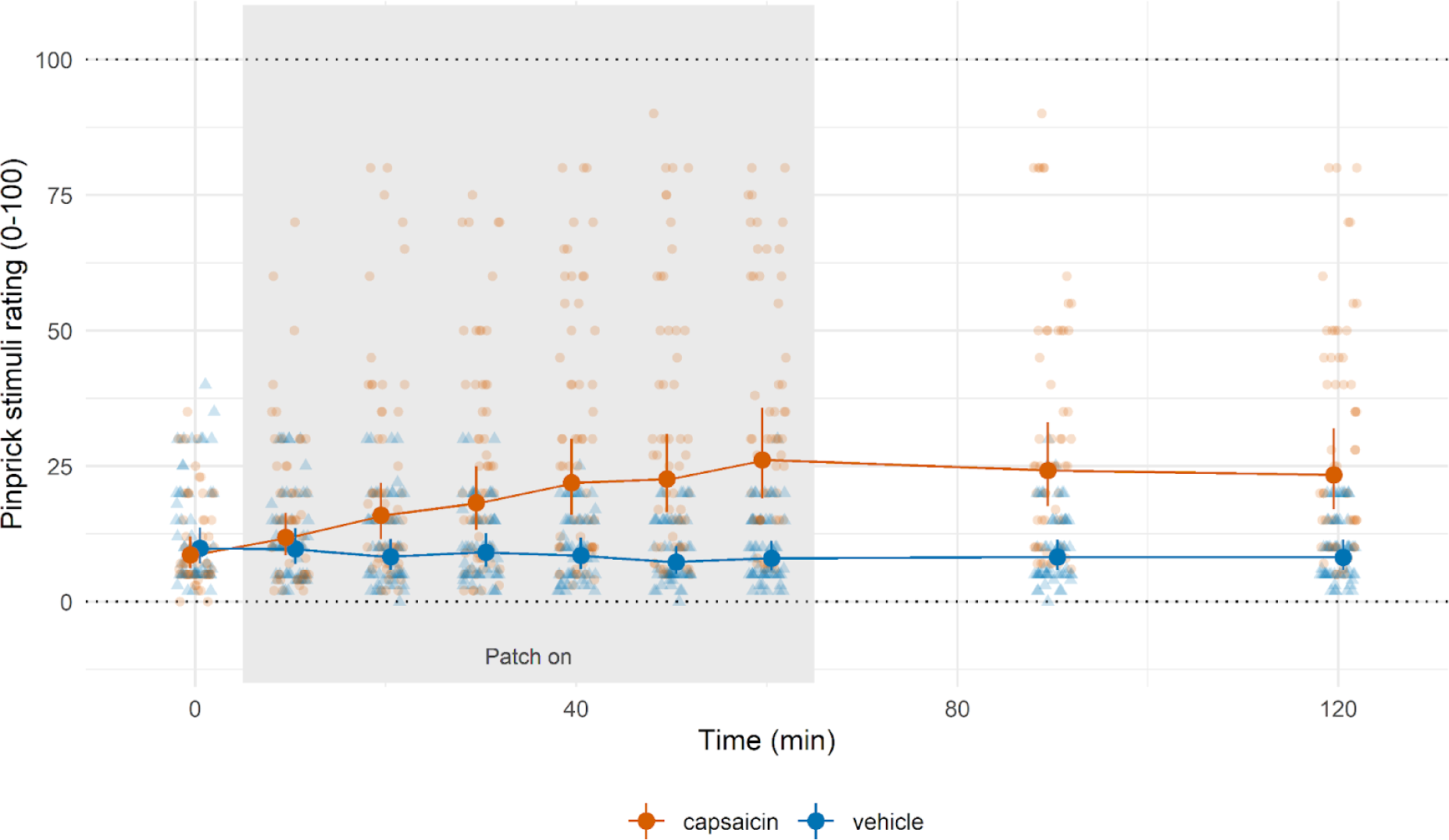
Perceived intensity of the pinprick stimuli, over time and across conditions. Individual ratings are represented as translucent dots in the background. The solid dots and vertical lines represent the marginal means and 95% confidence intervals fitted by the model. The solid lines connect the marginal means.

### Pinprick stimuli

‘Pinprick stimuli were mostly described as “pricking” and sometimes as “touch” (mostly for the early time points or for the vehicle arm, Figure 5).

Pinprick stimuli delivered on the forearm that received the vehicle patch (Figure 4, blue line) were perceived with a similar (or even slightly decreasing) intensity over time. Conversely, those delivered on the forearm receiving the capsaicin patch (Figure 4, dark-orange line) significantly increased over time and plateaued after patch removal, revealing the development of capsaicin-driven secondary mechanical hyperalgesia (Figure 4, Table 2). These changes in perceived intensity were accompanied by congruent changes in the quality of the sensations elicited by the pinprick stimuli (Figure 5, Table 3, 4).

**Figure 5:**
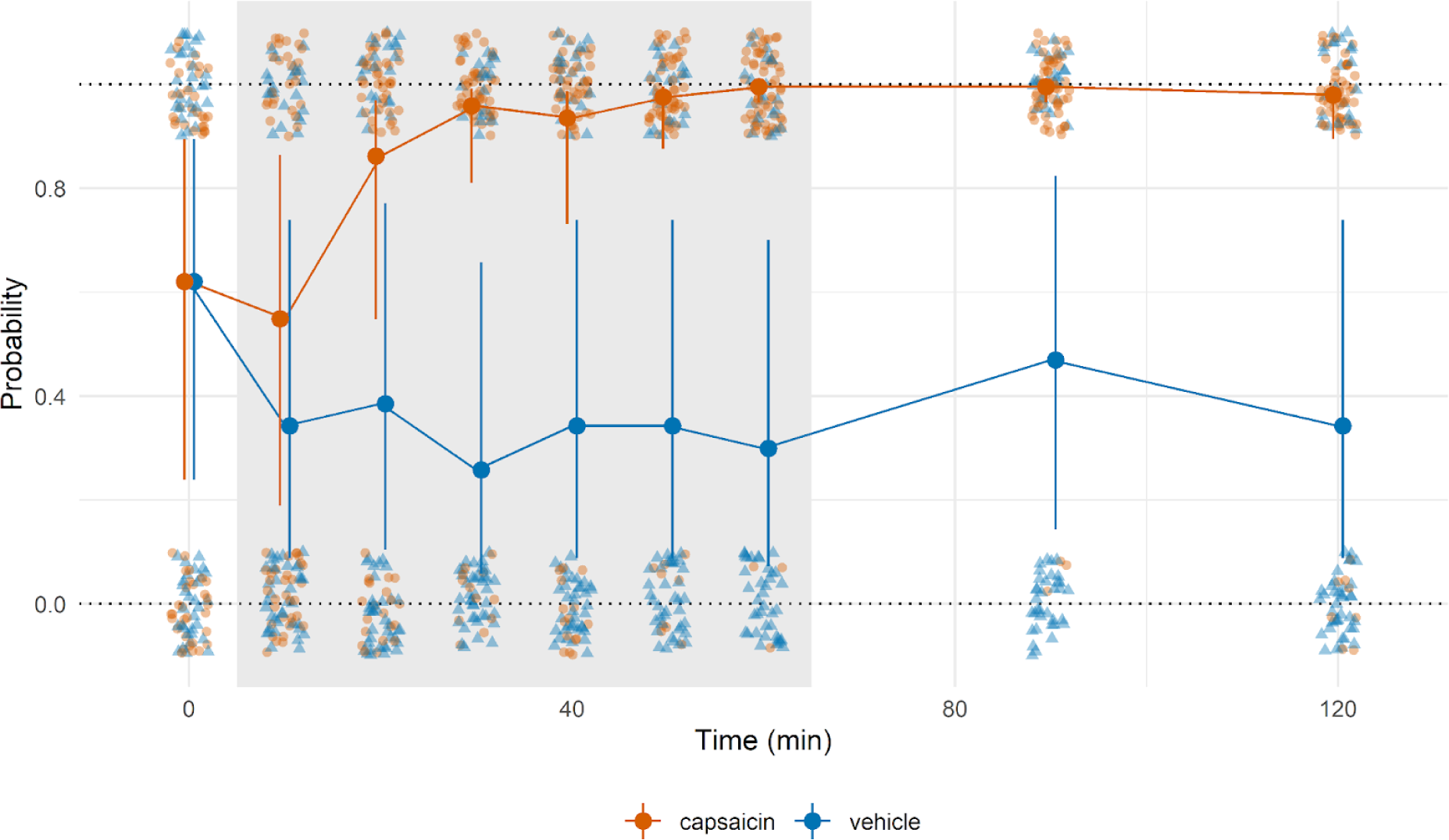
Qualitative assessment of sensations elicited by pinprick stimuli: probability of reporting a nociceptive descriptor (burning/pricking/painful). Individual responses are represented as translucent jittered dots in the background. The solid dots and vertical lines represent the marginal means and 95% confidence intervals fitted by the model. The solid lines connect the marginal means.

**Table 3:** Results of the pinprick ratings ANOVA.

| Factor | Sum Square | Mean Square | Num DF | Den DF | F value | Pr(>F) |
| --- | --- | --- | --- | --- | --- | --- |
| Time point | 18.253 | 2.282 | 8 | 990 | 13.748 | <0.001 |
| patch | 126.583 | 126.583 | 1 | 990 | 762.739 | <0.001 |
| Time point: patch | 40.797 | 5.100 | 8 | 990 | 30.729 | <0.001 |

**Table 4:** Results of the pinprick quality ANOVA.

| Factor | Chi Square | DF | Pr(>ChSq) |
| --- | --- | --- | --- |
| Time point | 93.221 | 8 | <0.001 |
| patch | 0.000 | 1 | 1.000 |
| Time point: patch | 73.512 | 8 | <0.001 |

### Temporal association

As can be seen in Figure 6, the arm raising effect (on capsaicin patch ratings) and mechanical secondary hyperalgesia seemed to largely follow the same time course, which was very different from that of the capsaicin patch sensations recorded in the horizontal resting position. However, the last set of measurements revealed that the arm raising effect and secondary hyperalgesia time courses were in fact distinct, as they dramatically diverged at this later time point. Indeed, whereas virtually no difference could be observed between the +90 and +120 time points for pinprick hyperalgesia, the arm raising effect, which was near maximal at 90 minutes post patch application, had (almost) completely disappeared at +120 minutes.

**Figure 6:**
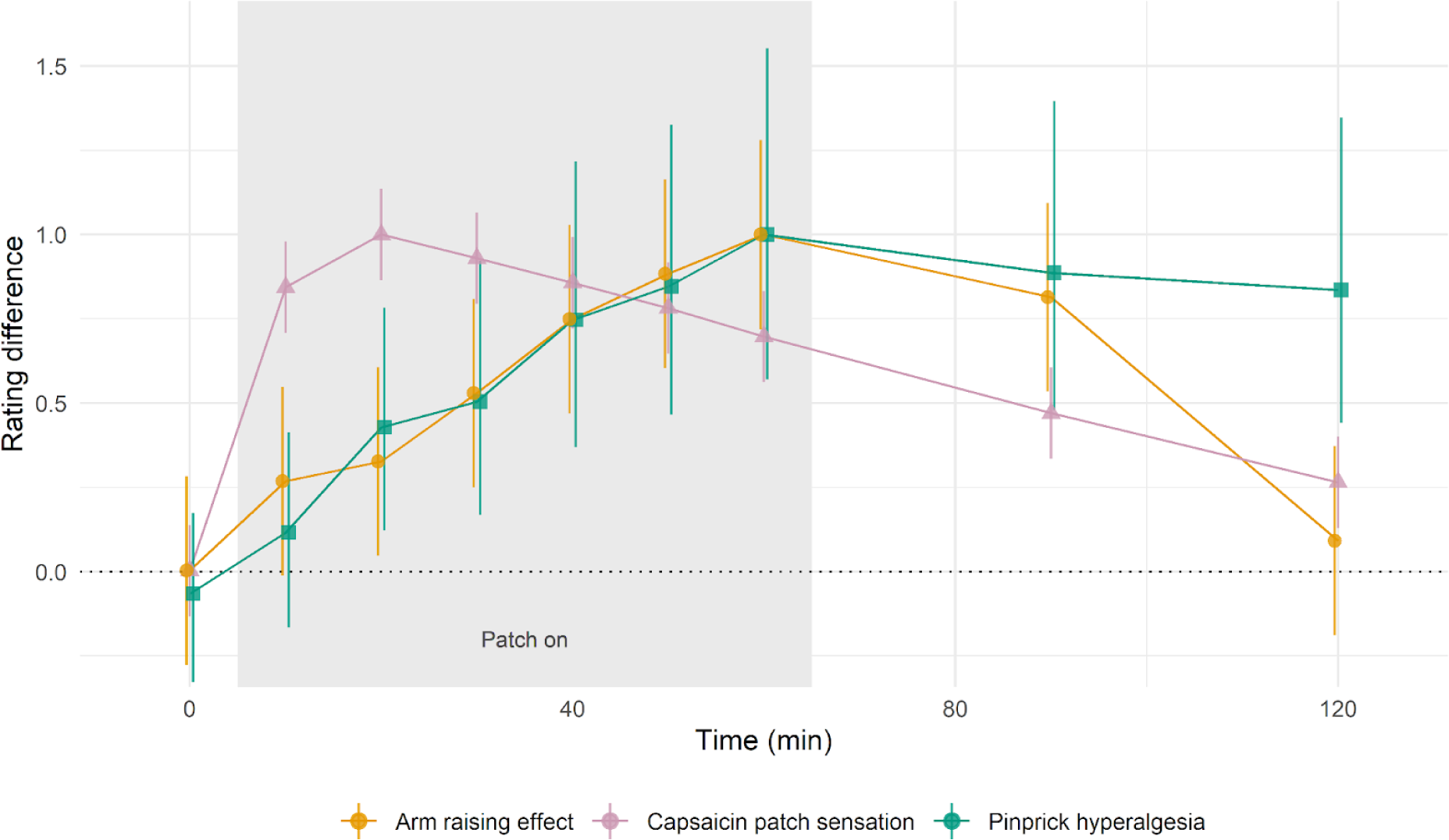
Normalized temporal evolution of the capsaicin patch sensation, the arm raising effect on the capsaicin patch sensation, and the capsaicin-driven mechanical hyperalgesia. The solid dots and vertical lines represent the marginal means and 95% confidence obtained from the previously described models and normalized so that the maximal marginal mean for each time course is equal to one. Solid lines connect the marginal means. The time course of the “capsaicin patch sensation” (pink) corresponds to the marginal mean for capsaicin patch ratings with the arm at rest (horizontal position), regardless of body position (shown not to affect these ratings). The “arm raising effect” time course (orange) corresponds to the marginal difference between capsaicin patch ratings obtained in the two arm positions (vertical – horizontal), regardless of body position. The “pinprick hyperalgesia” time course (green) corresponds to the marginal difference between intensity ratings of pinprick stimuli delivered on the two forearms (capsaicin – vehicle). For ease of comparison, the time courses were normalized by dividing (for each variable separately) by the largest marginal mean, so that each time course saturated at 1.

## 4. Discussion

The present data confirm that raising the arm during topical capsaicin treatment leads to a significant increase of the pain sensations experienced at the site of the patch (i.e. the arm raising effect). This effect did not appear to be mediated by the position of the arm relative to the body but rather by its position relative to the ground and is therefore likely caused by gravity. In parallel, as expected [28,32], we also observed the development of an increased sensitivity to pinprick stimuli (mechanical secondary hyperalgesia) in the skin area adjacent to the capsaicin patch. Inspection of their temporal evolution revealed that the development of the burning pain evoked by capsaicin, the development of the arm raising effect, and the development of secondary mechanical hyperalgesia followed three distinct time courses.

### The arm raising effect is likely caused by gravity-induced changes in the vascular system

Raising the arm from a horizontal to a vertical position and maintaining that position involves changes in the activity of various muscles of the arm and back. However, it seems unlikely that these changes are causing the arm raising effect as the muscles which would be the most involved in holding the arm vertically (*triceps brachialis* for elbow extension, the muscles involved in shoulder flexion and stabilization) are not located close to the patch [17]. Similarly, the muscles situated directly below the skin of the anterior forearm are mostly involved in mobilization of the wrist/hand and do not seem to be particularly involved in maintaining the arm in raised position [17].

Furthermore, unpublished data collected during piloting show that there is no difference in pain ratings when participants actively raise their arm vertically compared to when their arm is held in that position passively by the experimenter. Even though we cannot completely rule out muscular activity in the passive mobilization condition (the reason why this manipulation was abandoned in the final study protocol) this experimental manipulation should have led to more muscular activity in the active than the passive condition and, therefore, to changes in the pain ratings if the arm raising effect is indeed driven by muscular activity. This appeared not to be the case.

A more likely explanation is that the arm raising effect is related to changes in the vascular system. When the arm is raised, a large proportion of the blood contained in that limb is drained by gravity, superficial veins collapse and the local blood flow/blood pressure suddenly drops [30]. Given the relatively modest volume of blood contained in a single upper limb, this is not sufficient to activate baroreceptors and trigger a systemic response [31]. Instead, the preservation of perfusion relies primarily on the veno-arteriolar reflex, a local mechanism countering changes in transmural pressure by adapting the degree of vasodilation/vasoconstriction of the blood vessels (in this case increasing vasodilation in response to a drop of pressure)[31]. The fact that reducing blood flow by means of tourniquet constriction, rather than gravity, also leads to a sudden increase in the pain caused by capsaicin injection seems to corroborate this explanation of our observation [10].

### The arm raising effect is not another manifestation of the heterosynaptic LTP reflected by pinprick hypersensitivity at the skin

Interestingly, the development of the arm raising effect and of mechanical hyperalgesia appeared to be strongly correlated for the first hour (when the patch was on the skin), suggesting that the strong and sustained activation of epidermal nociceptors sensitized by capsaicin is probably a common driving factor of both phenomena. This initial close association is reminiscent of the observation by Byas-Smith *et al.* that tourniquet vascular occlusion led to an increase of capsaicin evoked pain only after pinprick hyperalgesia had developed [10].

As secondary hyperalgesia is thought to reflect heterosynaptic LTP in the dorsal horn nociceptive pathways, Byas-Smith’s observation led us to postulate that the arm raising effect could be another perceptual manifestation of this mechanism, amplifying the input of vascular sensory fibers in addition to that of mechanosensitive epidermal nociceptors [9,18]. This experiment was designed to test this hypothesis. The strong dissociation of time courses between the arm-raising effect (completely gone 60’ after patch removal) and secondary hyperalgesia at the skin (almost no decrease between patch removal and 60’ later) at later time points seems to rule out the possibility of a shared mechanism.

### Speculative mechanisms of the arm raising effect

One explanation of the arm raising effect could be that the decreased blood flow during arm raising leads to changes in the concentration of chemicals in the skin/perivascular milieu which would in turn modulate the activity of nociceptors and drive the increase in pain. For example, reduced blood flow could lead to a reduced clearance and therefore accumulation of chemical compounds released in reaction to capsaicin application to the skin. Alternatively, it could lead to a local hypoxia to which certain cells react by releasing vasodilatory chemicals which could also be pronociceptive, such as nitric oxide [19,30]. However, such mechanisms relying on diffusional changes in chemical concentration in the extracellular milieu are bound to be rather slow and therefore cannot account for the large and quick pain increase observed upon arm elevation (sometimes within less than 2 s) and the similarly fast decrease in pain perception upon return to the horizontal resting position.

Another explanation could rely on the peripheral sensitization of nociceptive fibers innervating the perivascular space or blood vessels, which could be able to respond to the vascular changes caused by the arm raising procedure (*e.g.* vascular wall stretch) and therefore cause the increase in pain directly. Arndt and Klement have described polymodal nociceptors innervating vein walls which were sensitive to stretch, temperature, osmotic pressure and electrical stimulations [5,22,23]. It has also been shown that the presence of capsaicin in the perivascular space leads to pain [4]. More recently, it has been suggested that not all of the small fibers innervating the arteriole-venule shunts are sympathetic but that some could be nociceptors [2,7,34]. Such nociceptors could be sensitized directly by capsaicin. In this case the delayed onset of the arm raising effect would be explained by the diffusion time necessary for capsaicin to reach deeper skin layers. The disappearance of the arm raising effect before spontaneous capsaicin pain completely fades away could similarly be attributed to a faster capsaicin clearance in the vicinity of the blood vessels [6,13].

Alternatively, sensitization of these nerve fibers could also be caused by other chemicals released in reaction to the topical application of capsaicin [8]. However, to be compatible with the temporal evolution of the arm raising effect (i.e. the delayed onset of the arm-raising effect in relation to the spontaneous burning sensation and its steady increase over time plateauing at 20 minutes after patch application), the concentration and effect of these compounds would need to steadily increase until patch removal, a pattern which cannot be ruled out, but does not appear likely.

One could also imagine that sensitization of vascular innervation (sensory and/or sympathetic) does not directly gives rise to the increased pain but does so by modulating the gating of nociceptive information coming from vascular and/or epidermal nociceptors (i.e. viscero-somatic convergence) [21,35].

Finally, the involvement of vascular nociceptors is perhaps supported by the fact that, in the experience of the authors, the pain when the arm was raised felt deeper and more widespread than when the arm was at rest. This pain also did not seem to habituate and the quality of the pain sensation was different in that it had a sharp, stabbing and pulsatile quality that the “spontaneous” capsaicin pain did not have. These observations are reminiscent of the description of the pain triggered directly by vascular stimulation made by Arndt & Klement [5].

Several of the aforementioned mechanisms rely on peripheral sensitization of nerve fibers. Interestingly, several non-neural cells, like smooth muscle cells, also express TRPV1 which has been linked to the rapid regulation of myogenic tone [20,33]. In the skin, non-neural cells like Schwann cells and keratinocytes have been shown to contribute to the detection of noxious stimuli and pain sensation [1,15,29]. It therefore seems possible that vascular smooth muscle cells or endothelial cells themselves could contribute to peripheral sensitization.

## 5. Conclusion

In this study, we showed that vascular changes (caused by a change of the upper limb position) can dramatically modulate the pain sensations caused by the topical application of capsaicin. Even though the exact mechanism underlying this increase in pain sensation is not entirely clear, it seems to involve nerve fibers innervating blood vessels. Knowledge of the sensory innervation of blood vessels and of possible crosstalks between the autonomic and nociceptive systems remains limited. In the future, the phenomenon described in this paper could prove a useful experimental model to investigate these topics.

## References

[1] Abdo H, Calvo-Enrique L, Lopez JM, Song J, Zhang MD, Usoskin D, El Manira A, Adameyko I, Hjerling-Leffler J, Ernfors P. Specialized cutaneous Schwann cells initiate pain sensation. Science (New York, NY) 2019;365:695–699.

[2] Albrecht PJ, Hou Q, Argoff CE, Storey JR, Wymer JP, Rice FL. Excessive Peptidergic Sensory Innervation of Cutaneous Arteriole–Venule Shunts (AVS) in the Palmar Glabrous Skin of Fibromyalgia Patients: Implications for Widespread Deep Tissue Pain and Fatigue. Pain Medicine 2013;14:895–915.

[3] Allaire J. RStudio: integrated development environment for R. Boston, MA 2012;770:165–171.

[4] Arndt JO, Kindgen-Milles D, Klement W. Capsaicin did not evoke pain from human hand vein segments but did so after injections into the paravascular tissue. The Journal of Physiology 1993;463:491–499.

[5] Arndt JO, Klement W. Pain evoked by polymodal stimulation of hand veins in humans. J Physiol 1991;440:467–478.

[6] Babbar S, Marier J-F, Mouksassi M-S, Beliveau M, Vanhove GF, Chanda S, Bley K. Pharmacokinetic Analysis of Capsaicin After Topical Administration of a High-Concentration Capsaicin Patch to Patients With Peripheral Neuropathic Pain. Therapeutic Drug Monitoring 2009;31:502.

[7] Bowsher D, Woods GC, Nicholas AK, Carvalho OM, Haggett CE, Tedman B, Mackenzie JM, Crooks D, Mahmood N, Twomey AJ, Hann S, Jones D, Wymer JP, Albrecht PJ, Argoff CE, Rice FL. Absence of pain with hyperhidrosis: A new syndrome where vascular afferents may mediate cutaneous sensation. PAIN 2009;147:287.

[8] Braga Ferreira LG, Faria JV, Dos Santos JPS, Faria RX. Capsaicin: TRPV1-independent mechanisms and novel therapeutic possibilities. European Journal of Pharmacology 2020;887:173356.

[9] van den Broeke EN. Central sensitization and pain hypersensitivity: Some critical considerations. F1000Research 2018;7:1325.

[10] Byas-Smith MG, Bennett GJ, Gracely RH, Max MB, Robinovitz E, Dubner R. Tourniquet constriction exacerbates hyperalgesia-related pain induced by intradermal capsaicin injection. Anesthesiology 1999;91:617–25.

[11] Carpenter SE, Lynn B. Vascular and sensory responses of human skin to mild injury after topical treatment with capsaicin. Br J Pharmacol 1981;73:755–758.

[12] Caterina MJ, Pang Z. TRP Channels in Skin Biology and Pathophysiology. Pharmaceuticals (Basel, Switzerland) 2016;9.

[13] Chanda S, Bashir M, Babbar S, Koganti A, Bley K. In Vitro Hepatic and Skin Metabolism of Capsaicin. Drug Metab Dispos 2008;36:670–675.

[14] Courtin AS, Mouraux A. Combining Topical Agonists With the Recording of Event-Related Brain Potentials to Probe the Functional Involvement of TRPM8, TRPA1 and TRPV1 in Heat and Cold Transduction in the Human Skin. The journal of pain : official journal of the American Pain Society 2022;23:754–771.

[15] Denda M, Tsutsumi M. Roles of transient receptor potential proteins (TRPs) in epidermal keratinocytes. Advances in experimental medicine and biology 2011;704:847–60.

[16] Foreman JC, Jordan CC, Oehme P, Renner H. Structure-activity relationships for some substance P-related peptides that cause wheal and flare reactions in human skin. J Physiol 1983;335:449–465.

[17] Hamill J, Knutzen KM. Biomechanical Basis of Human Movement. Lippincott Williams & Wilkins, 2006.

[18] Henrich F, Magerl W, Klein T, Greffrath W, Treede R-D. Capsaicin-sensitive C- and A-fibre nociceptors control long-term potentiation-like pain amplification in humans. Brain 2015;138:2505–2520.

[19] Holthusen H, Arndt JO. Nitric oxide evokes pain in humans on intracutaneous injection. Neuroscience letters 1994;165:71–4.

[20] Jackson WF. The heat is on! TRPV1 channels and resistance artery myogenic tone. J Physiol 2022;600:2021–2022.

[21] Jänig W. Neurobiologie viszeraler Schmerzen. Schmerz 2014;28:233–251.

[22] Klement W, Arndt JO. Pain but no temperature sensations are evoked by thermal stimulation of cutaneous veins in man. Neuroscience letters 1991;123:119–22.

[23] Klement W, Arndt JO. The role of nociceptors of cutaneous veins in the mediation of cold pain in man. The Journal of physiology 1992;449:73–83.

[24] Kuznetsova A, Brockhoff PB, Christensen RHB. lmerTest Package: Tests in Linear Mixed Effects Models. J Stat Soft 2017;82:1–26.

[25] Lüdecke D. ggeffects: Tidy Data Frames of Marginal Effects from Regression Models. Journal of Open Source Software 2018;3:772.

[26] Moisset X, Bouhassira D, Attal N. French guidelines for neuropathic pain: An update and commentary. Revue Neurologique 2021;177:834.

[27] van Neerven SGA, Mouraux A. Capsaicin-Induced Skin Desensitization Differentially Affects A-Delta and C-Fiber-Mediated Heat Sensitivity. Front Pharmacol 2020;11:615.

[28] O’Neill J, Brock C, Olesen AE, Andresen T, Nilsson M, Dickenson AH. Unravelling the Mystery of Capsaicin: A Tool to Understand and Treat Pain. Pharmacol Rev 2012;64:939–971.

[29] Pang Z, Sakamoto T, Tiwari V, Kim YS, Yang F, Dong X, Guler AD, Guan Y, Caterina MJ. Selective keratinocyte stimulation is sufficient to evoke nociception in mice. Pain 2015;156:656–65.

[30] Pappano AJ, Gil Wier W. 9 - The Peripheral Circulation and its Control. In: Pappano AJ, Gil Wier W, editors. Cardiovascular Physiology (Tenth Edition). Philadelphia: Elsevier, 2013. pp. 171–194. doi:10.1016/B978-0-323-08697-4.00009-5.

[31] Low PA. 38 - Venoarteriolar Reflex. In: Robertson D, Biaggioni I, Burnstock G, Low PA, editors. Primer on the Autonomic Nervous System (Second Edition). San Diego: Academic Press, 2004. pp. 152–153. doi:10.1016/B978-012589762-4/50039-6.

[32] Petersen KL, Rowbotham MC. A new human experimental pain model: the heat/capsaicin sensitization model. NeuroReport 1999;10:1511.

[33] Phan TX, Ton HT, Gulyás H, Pórszász R, Tóth A, Russo R, Kay MW, Sahibzada N, Ahern GP. TRPV1 in arteries enables a rapid myogenic tone. The Journal of Physiology 2022;600:1651–1666.

[34] Rice FL, Albrecht PJ. 6.01 - Cutaneous Mechanisms of Tactile Perception: Morphological and Chemical Organization of the Innervation to the Skin. In: Masland RH, Albright TD, Albright TD, Masland RH, Dallos P, Oertel D, Firestein S, Beauchamp GK, Catherine Bushnell M, Basbaum AI, Kaas JH, Gardner EP, editors. The Senses: A Comprehensive Reference. New York: Academic Press, 2008. pp. 1–31. doi:10.1016/B978-012370880-9.00340-6.

[35] Schwartz ES, Gebhart GF. Visceral Pain. In: Taylor BK, Finn DP, editors. Behavioral Neurobiology of Chronic Pain. Current Topics in Behavioral Neurosciences. Berlin, Heidelberg: Springer, 2014. pp. 171–197. doi:10.1007/7854_2014_315.

[36] Sultana A, Singla RK, He X, Sun Y, Alam MS, Shen B. Topical Capsaicin for the Treatment of Neuropathic Pain. Current Drug Metabolism n.d.;22:198–207.

[37] Voets T, Droogmans G, Wissenbach U, Janssens A, Flockerzi V, Nilius B. The principle of temperature-dependent gating in cold- and heat-sensitive TRP channels. Nature 2004;430:748–54.

[38] Wickham H, Averick M, Bryan J, Chang W, McGowan LD, François R, Grolemund G, Hayes A, Henry L, Hester J. Welcome to the Tidyverse. Journal of open source software 2019;4:1686.

